# A generalized metabolic theory for marine ecosystems

**DOI:** 10.64898/2026.07.11.737976

**Authors:** Andrea Tabi

## Abstract

Marine species inhabit a three-dimensional stratified environment where metabolism is simultaneously influenced by decreasing temperature and increasing hydrostatic pressure with ocean depth. While the Metabolic Theory of Ecology (MTE) incorporates the effects of body mass and temperature, for marine organisms it neglects one of the fundamental thermodynamic state variables governing biochemical reaction kinetics: hydrostatic pressure. We derive a generalized metabolic theory by extending MTE with transition-state theory by explicitly incorporating pressure sensitivity into the Arrhenius formulation. The resulting model predicts that hydrostatic pressure increases the effective activation energy of biochemical reactions, leading to progressively lower metabolic rates with increasing depth. We tested this prediction using a global database comprising 689 metabolic measurements across 11 marine phyla and 22 taxonomic classes. Incorporating hydrostatic pressure substantially improved model performance relative to the classical MTE. Across major marine taxa, activation energy and activation volume varied largely independently, suggesting that pressure adaptation does not require corresponding changes in thermal sensitivity. Our results suggest that hydrostatic pressure is a fundamental thermodynamic constraint that regulates the pace of life across the ocean and consequently the ecological dynamics across Earth’s largest biome.

## I. INTRODUCTION

Oceans provide the only major biome where organisms routinely experience vertical gradients in both hydrostatic pressure and temperature. These environmental axes fundamentally influence physiological performance, species distributions, community assembly and biodiversity patterns across the ocean depth [16, 17]. Understanding how temperature and pressure jointly regulate organismal energetics is therefore essential for improving conservation and management planning of marine biodiversity under global environmental change [14].

Metabolic theory of ecology (MTE) is one of the most prominent ecological theories, linking individual energetics to fundamental biological properties such as body mass and environmental temperature [10, 25]. These scaling relationships have provided the basis for allometric scaling models capable of making predictions from individuals to communities and ecosystems [1, 41, 49]. While the mechanistic origins of allometric scaling remain debated [3, 20, 26, 29, 38, 47, 49], the temperature dependence of metabolic rates is well established from chemical thermodynamics. Harlow Shapley first demonstrated the temperature dependence of ant locomotion, which then contributed to the subsequent formulation of the temperature dependence of biochemical reaction rates using Arrhenius–Boltzmann kinetics [35]. Ever since, the Arrhenius equation has been widely used as an approximation for temperature-dependent biological rates in ecological and trait-based models [10, 19, 41]. However, biochemical reaction rates depend not only on temperature [40]. Transition-state theory, formalized by the Eyring equation, provides a more general description of chemical kinetics by assuming that every reaction passes through a short, high-energy intermediate known as the transition state or activated complex [23, 32]. Importantly, the free-energy barrier associated with this transition state depends on both temperature and pressure through the activation energy and activation volume. Therefore, the classical MTE can be viewed as the low-pressure approximation of a more general metabolic theory. For terrestrial organisms, and for shallow-water systems, hydrostatic pressure differences are negligible, however marine organisms inhabit environments in enormous depth gradients, where hydrostatic pressure exerts substantial physiological effects. Previous studies have shown large differences in respiration rates between shallow- and deep-water species after temperature and body size normalisation, suggesting an important role for pressure adaptation [6–9, 24, 42]. Marine organisms in bathyal, abyssal and hadal environments are exposed to high hydrostatic pressures at nearly constant low temperatures (Fig. 1a). High pressure perturbs protein structure, membrane fluidity and enzyme kinetics, reducing aerobic performance unless compensated by evolutionary adaptations. Deep-sea organisms exhibit numerous biochemical adaptations, including modified metabolic enzymes, altered membrane lipid composition, accumulation of trimethylamine N-oxide (TMAO), cytoskeletal modifications and molecular chaperones that partially restore cellular function under pressure[8, 24]. Nevertheless, even after accounting for body size and temperature differences, deep-sea species typically exhibit substantially lower metabolic rates than their shallow-water relatives regardless of food and oxygen availability. This suggests that hydrostatic pressure together with low temperatures fundamentally alter metabolism setting physiological constraints to the pace of life [6, 52].

**Figure 1.**
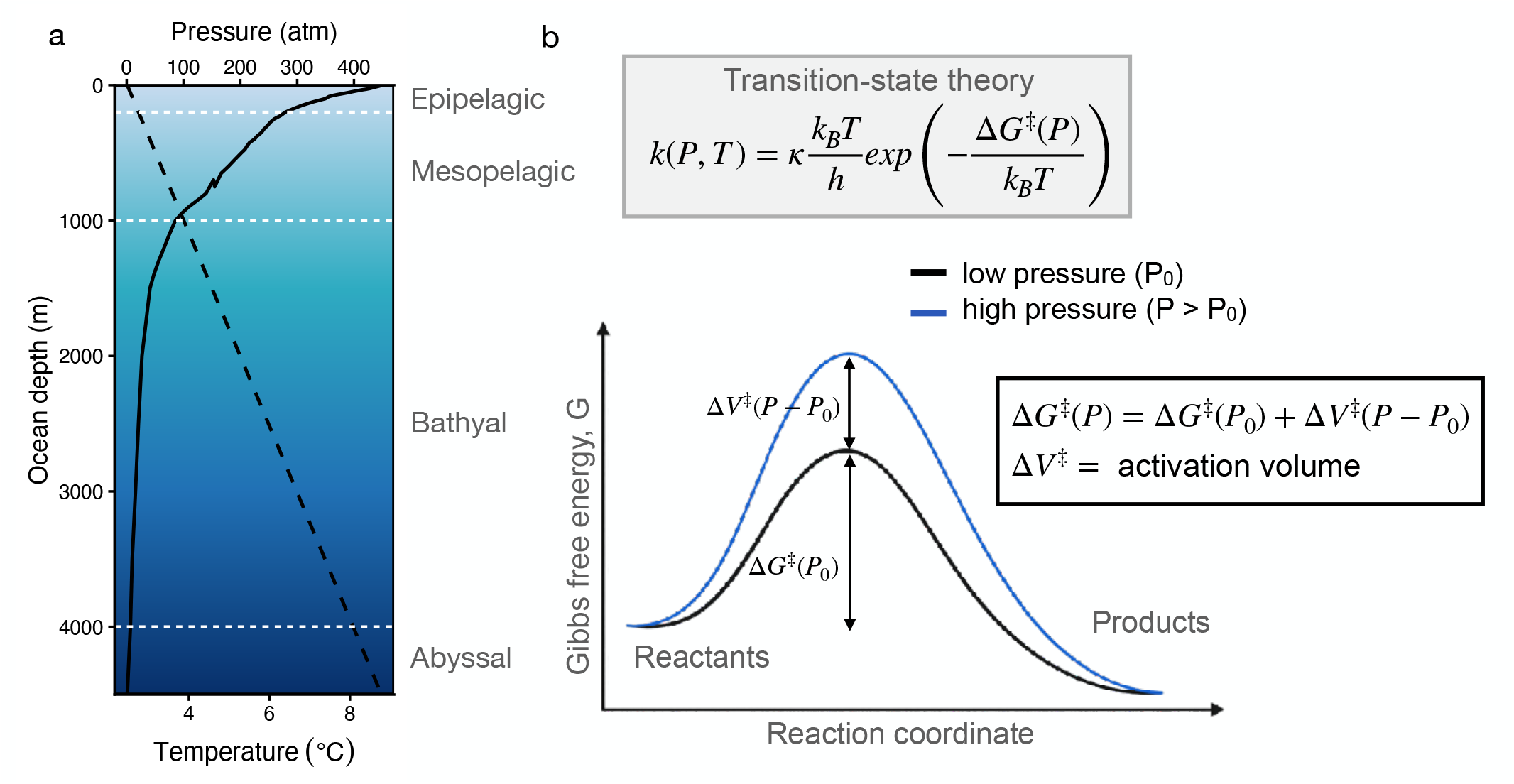
Temperature and hydrostatic pressure across the ocean depth and their mechanistic effects on biochemical reaction rates. (a) Mean ocean temperature decreases rapidly and nonlinearly, whereas hydrostatic pressure increases linearly across ocean depth. Dashed lines indicate the approximate boundaries between the epipelagic, mesopelagic, bathyal and abyssal zones. Dashed line: pressure, solid line:temperature. (b) Transition-state theory predicts that both temperature and hydrostatic pressure change biochemical reaction rates. Hydrostatic pressure increases the activation free energy of biochemical reactions, thereby slowing reaction rates. This provides the mechanistic basis for incorporating hydrostatic pressure into metabolic scaling theory.

**Figure 2.**
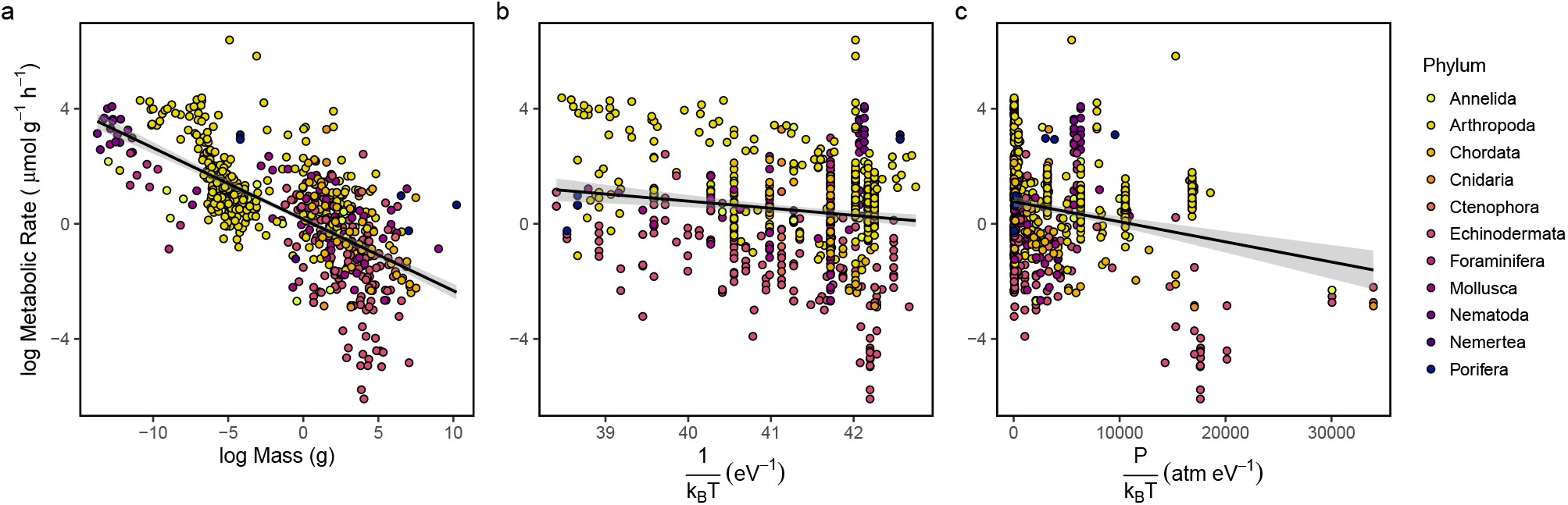
Relationships between mass-specific metabolic rate and (a) body mass, (b) temperature term (1*/k*_*B*_ *T*), and (c) the pressure-corrected temperature term (*P/k*_*B*_ *T*). Points are colored by phyla in each panel.

**Figure 3.**
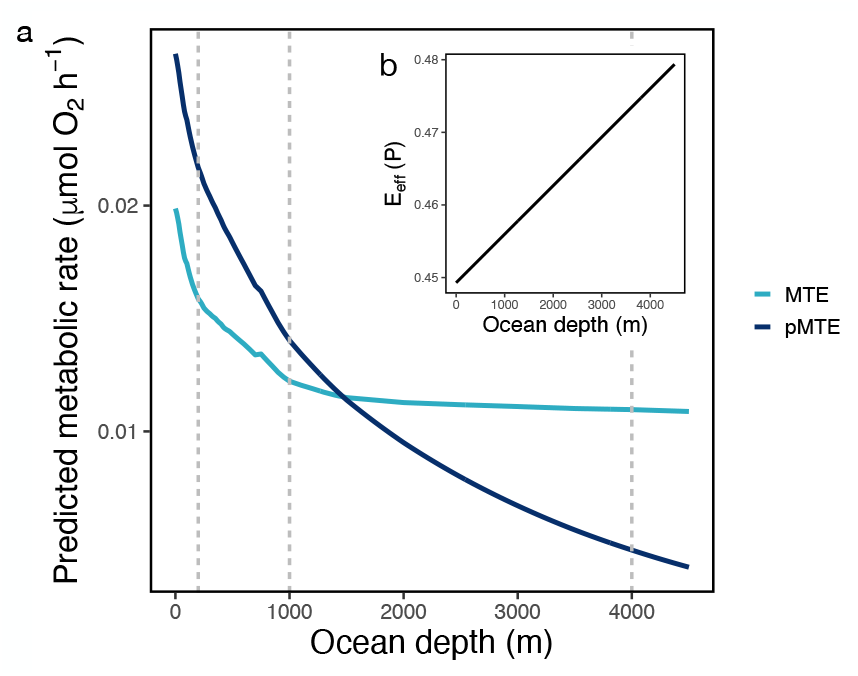
Predicted metabolic rates across ocean depth under classical metabolic theory (MTE) and the pressure-corrected model (pMTE). (a) Predictions are shown for a 10 g organism using the observed ocean temperature profile. Vertical dashed lines indicate the approximate boundaries between the epipelagic (0-200 m), mesopelagic (200-1000 m), bathyal (1000-4000 m) and abyssal (>4000m) zones. (b) Pressure linearly increases the effective activation energy across the depth gradient.

**Figure 4.**
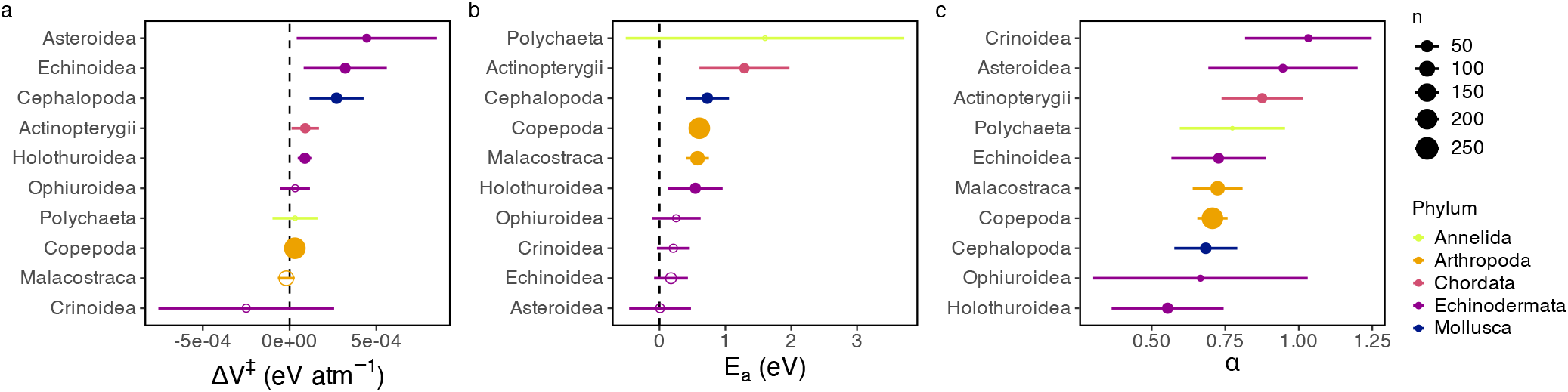
Estimated metabolic scaling components by marine taxa. Class-specific estimates of activation volume (Δ*V* ^*‡*^), activation energy (*E*_*a*_), and mass-scaling exponent (*α*) for taxonomic classes with *n* > 10. Points indicate maximumlikelihood estimates, horizontal bars denote 95% confidence intervals, point size is proportional to sample size, and colors indicate phylum. Parameters were estimated separately for each major marine taxonomic class. Activation volume varied independently of both activation energy and the mass-scaling exponent, indicating that pressure sensitivity represents an additional axis of physiological variation rather than a simple consequence of thermal sensitivity or body-size scaling.

Although recent reviews have argued that transition-state theory provides a more appropriate description of enzyme kinetics than the Arrhenius approximation used in metabolic theory [13], the implications of transition-state theory for metabolic scaling have not been developed. In particular, the pressure dependence of the activation free energy—and the resulting activation volume term—has not been incorporated into the metabolic theory of ecology. Here, we present a generalized formulation of the Metabolic Theory of Ecology that explicitly incorporates both thermal and pressure sensitivity based on transition-state theory. We then test this framework using a global dataset of marine metabolic rates and show that including hydrostatic pressure significantly improves predictions of metabolic scaling across marine taxa.

## II. PRESSURE-CORRECTED METABOLIC THEORY OF ECOLOGY

The MTE assumes that metabolic rates are governed by body mass and temperature according to the Arrhenius kinetics:

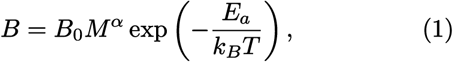

where *B* is metabolic rate, *M* is body mass, *α* is the metabolic scaling exponent, *E*_*a*_ is the activation energy, *k*_*B*_ is Boltzmann’s constant, and *T* is environmental temperature. Equation (1) implicitly assumes constant pressure. While this assumption is appropriate for terrestrial systems, hydrostatic pressure increases from approximately 1 atmosphere at the ocean surface to more than 1000 atmospheres in the deepest marine trenches. Therefore, hydrostatic pressure is expected to modify biochemical reaction rates.

To account for this effect, we apply the Eyring-Polányi equation [22, 23] in line with the transition-state theory [32], where the reaction rate is expressed as

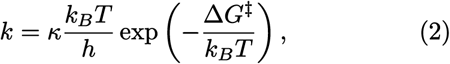

where *κ* is the transmission coefficient, *h* is Planck’s constant, and Δ*G*^‡^ is the Gibbs free energy of activation. The Gibbs free energy depends on pressure according to *dG* = *V dP* − *S dT*, where *V* is the volume, *P* is the pressure, *T* is the temperature and *S* is the entropy of the system. Thus, at constant temperature, the Gibbs free energy of activation depends only on the activation volume:

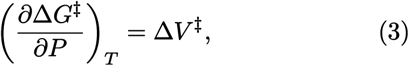

where Δ*V* ^‡^ is the activation volume, representing the volume difference between the transition state and the reactant state. Assuming that Δ*V* ^‡^ is approximately constant over the relevant pressure range, we can calculate the free energy of the system

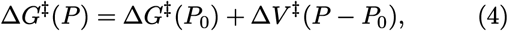

where *P*_0_ is a reference pressure. This gives the pressure-correction term due to volume changes during the chemical reaction (Fig.1B). If we substitute this into Eq. (2) we get the reaction rate

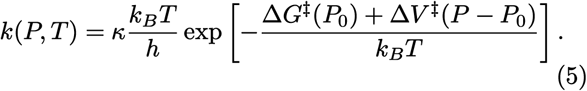

Assuming organismal metabolic rate is proportional to the underlying biochemical reaction rate, The prefactor is absorbed into the normalization constant 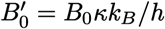, and we define *E*_*a*_ ≈ Δ*G*^‡^(*P*_0_). This gives

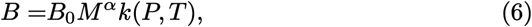

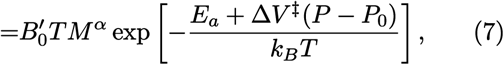

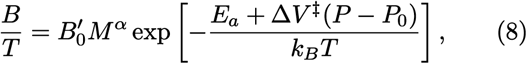

then taking logarithms,

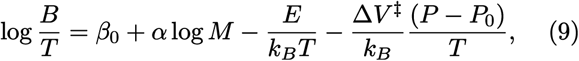

which predicts that metabolic rate depends linearly on body mass, the Arrhenius temperature term 1*/*(*k*_*B*_*T*), and the pressure-temperature interaction *P/*(*k*_*B*_*T*). The classical MTE is recovered as the special case, when Δ*V* ^‡^ = 0 corresponding to negligible pressure effects. Equation (7) predicts that temperature and pressure do not contribute independently to metabolism. Instead, hydrostatic pressure modifies the effective activation energy of biochemical reactions,

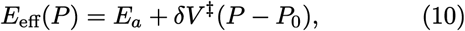

where (*E*_eff_) is the effective activation energy. Thus, increasing hydrostatic pressure increases the energetic barrier that must be overcome for biochemical reactions to proceed. Consequently, the influence of pressure becomes progressively stronger as temperature decreases because both variables act jointly through the denominator (*k*_*B*_*T*). This formulation therefore predicts that pressure and temperature are not independent environmental drivers of metabolism; rather, hydrostatic pressure amplifies the temperature dependence of metabolic rates by increasing the effective activation energy.

## III. EMPIRICAL VALIDATION

The pressure-corrected model makes two predictions. First, incorporating hydrostatic pressure should improve the prediction of metabolic rates relative to the classical MTE. Second, if activation energy and volume represent general biochemical properties, these quantities should be conserved across marine taxa. To test these predictions, we analysed the metabolic rates of marine species using a global dataset ranging across 11 phyla and 22 taxonomic classes (Fig.2). First, we contrasted the pressure-corrected metabolic model (M3) against the classical MTE (M2) and a body mass scaling model (M1) performed on whole-organism metabolic rates (*µ*mol h^−1^) using linear models (Table 1). Body mass explains a huge amount of variation (R^2^=0.89) with mass scaling exponent close to the canonical value of 3*/*4. The classical MTE model recovered an activation energy of *E*_*a*_ = 0.64 eV, close to the canonical value used in metabolic theory (0.6 −0.7 eV). However, when the pressure-temperature term was included, the estimated thermal activation energy decreased from 0.64 to 0.45 eV while revealing a strong negative pressure effect. Overall, the pressure-corrected model gave the best model fit with highest explained variance (R^2^=0.93) and higher predictive ability compared to the classic MTE model (ΔAIC = 66). The classical MTE model largely overestimates the marine metabolic rates with increasing depth (Fig.3). Classical MTE predicts that metabolism approaches a plateau once ocean temperatures stabilize below the thermocline, whereas the pressure-corrected model predicts continued metabolic decline with increasing hydrostatic pressure. Cross-validation using shallow-water training data demonstrated consistently better predictive ability for pMTE than classical MTE across increasing test depths (see details in SM).

**Table I.** Comparison of metabolic scaling models. Model 1 corresponds to mass scaling alone, Model 2 is the classical Metabolic Theory of Ecology (MTE), and Model 3 is the proposed pressure-corrected model (pMTE) derived from transition-state theory.

| Model | Description | Equation | $\alpha$ | $\beta_T$ | $\beta_P$ | $R^2$ | AIC |
| --- | --- | --- | --- | --- | --- | --- | --- |
| M1 | Mass scaling | $\log(B/T) = \beta_0 + \alpha \log(M)$ | 0.75 | – | – | 0.89 | 2228 |
| M2 | MTE | $\log(B/T) = \beta_0 + \alpha \log(M) + \frac{E_a}{k_B T}$ | 0.71 | –0.64 | – | 0.92 | 2021 |
| M3 | pMTE | $\log(B/T) = \beta_0 + \alpha \log(M) + \frac{E_a}{k_B T} + \frac{\Delta V^\ddagger P}{k_B T}$ | **0.71** | **–0.45** | **–6.72e<sup>–5</sup>** | **0.93** | **1955** |

Next, we analysed the temperature and pressure sensitivity to taxonomic variability (Table 2). By allowing the intercept (Mix1) to vary among classes significantly improved the model fit compared to the linear model (M3). However, class-specific temperature and pressure sensitivity coefficients moderately improved the model fit (Δ*AIC* ≈ 13), indicating that the temperature and pressure dependence of metabolic rates vary moderately across the major marine animal classes represented in the dataset. Overall, most variation among taxa was explained by baseline metabolic differences, although modest class-specific differences in thermal and pressure sensitivity were also detected. This does not support the assumption of a common temperature- and pressure-dependent activation term predicted by transition-state theory.

**Table II.**
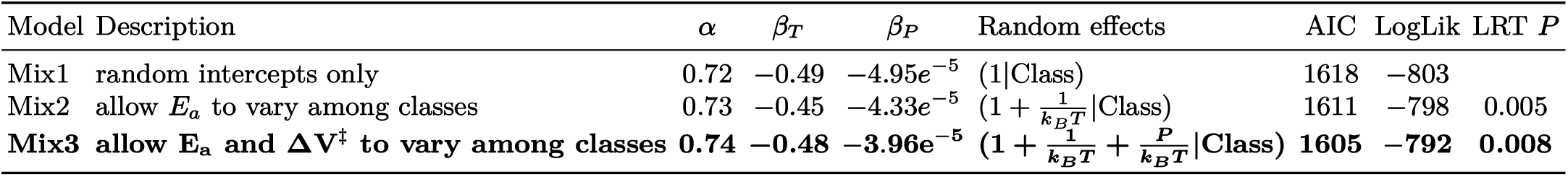
Comparison of mixed-effects pressure-corrected metabolic scaling models. All models include a random intercept for taxonomic class.

We quantified class-specific metabolic scaling components by separately fitting the pressure-corrected MTE models (Fig.4). All terms showed substantial variation across taxonomic classes. Activation energy and activation volume were not significantly correlated across taxonomic classes (Spearman’s *ρ* = −0.26, *P* = 0.47). This suggests that adaptation to high hydrostatic pressure does not necessarily require a corresponding change in thermal sensitivity. Activation energy and volume were also not strongly associated with mass-scaling exponent (Spearman’s *ρ* = −0.17, *P* = 0.63 and *ρ* = −0.03, *P* = 0.95).

## IV. DISCUSSION

Metabolic theory has become one of the fundamental theories linking organismal physiology to population, community and ecosystem processes through the effects of body size and temperature [1, 10, 41]. Our results suggest that extending this framework to marine ecosystems requires incorporating hydrostatic pressure as a fundamental thermodynamic variable governing biochemical reaction kinetics [13]. Rather than acting as an independent environmental driver, hydrostatic pressure modifies the effective activation energy of biochemical reactions through the activation volume predicted by transition-state theory. Therefore, the classical MTE is effectively a low-pressure approximation of a more general metabolic theory.

Classical MTE recovered the canonical scaling relationships of approximately 3*/*4 for body mass and 0.65 eV for activation energy, indicating that marine organisms broadly follow the same energetic constraints as terrestrial organisms [34]. However, our results demonstrate that metabolic rates in the ocean are more accurately described by the joint effects of body size, temperature and hydrostatic pressure. Note that the pressure–temperature interaction term was not introduced to improve statistical fit, but it follows directly from transition-state theory and therefore constitutes rather a mechanistic prediction.

We also detected significant variation among taxonomic classes in baseline metabolic rates and scaling relationships. These findings add to growing evidence that the parameters of metabolic scaling are not universal constants, but emergent properties shaped by physical constraints, intrinsic stochasticity, environmental conditions and evolutionary adaptation [26, 38, 47]. Together, these results support the view that metabolic scaling emerges from interacting biological and physical processes rather than fixed universal parameters.

The generalized metabolic theory has implications well beyond individual physiology because metabolic rates provide the energetic basis for population growth, species interactions, ecosystem respiration and biogeochemical cycling [1, 10, 41]. Classical MTE predicts that metabolic rates approach a plateau below the thermocline as ocean temperatures become nearly constant. In contrast, our theory predicts a continued decline in biological rates throughout the bathyal, abyssal and hadal zones due to progressively increasing hydrostatic pressure. This implies that ecological processes including growth, reproduction, competition, mortality, recovery and community turnover should also slow systematically with depth, providing a mechanistic connection between organismal energetics and the characteristically slow ecological dynamics of the deep ocean [17, 34, 42]. In other words, this framework provides a mechanistic basis for the hypothesis that ecological timescales increase systematically with ocean depth. While temperature largely determines the pace of life on land, temperature and hydrostatic pressure jointly determine the pace of life in the ocean. Pressure-induced reductions in metabolism are therefore expected to propagate across levels of biological organization, from individuals to ecosystems across the deep ocean. [6, 34, 44].

Our results also have important limitations. Pressure and temperature cannot be treated as statistically independent predictors because transition-state theory predicts that pressure modifies the activation free energy, producing the combined pressure-temperature term rather than an additive effect. Thus, our analysis quantifies the effective pressure dependence observed across marine taxa rather than separating immediate physiological responses from long-term evolutionary adaptation. Moreover, depth covaries with other environmental factors, including food supply, oxygen availability, carbonate chemistry and hydrographic conditions. Although previous studies generally report relatively weak effects of food availability and oxygen on metabolic rates after accounting for body size and temperature [6, 42], these effects remain incompletely understood, particularly in the deep sea. Future work should provide a more comprehensive theory of metabolic scaling across marine ecosystems including the role of additional environmental gradients and evolutionary adaptation.

Beyond its theoretical implications, the generalized metabolic theory has important consequences for the conservation management and risk assessment of marine ecosystems [18]. Deep-sea habitats are increasingly threatened by climate change and human activities such as bottom fishing and deep-seabed mining. Evidence show extremely slow recovery for abyssal ecosystems after disturbances caused by deep-sea mining [5, 30, 31, 43]. Yet ecological predictions for these environments remain largely based on metabolic models developed for terrestrial or shallow-water systems which systematically over-estimate metabolism and other biological rates. The generalised pressure-corrected metabolic model imposes slower growth, reproduction and population turnover throughout the deep ocean. Consequently, disturbed abyssal ecosystems are expected to exhibit longer recovery times and greater vulnerability to anthropogenic disturbance than predicted by classical metabolic theory. Incorporating hydrostatic pressure into ecological models should therefore improve predictions of ecosystem resilience, recovery and functioning across the full ocean depth gradient.

## V. METHODS

### A. Data

We compiled a global database of published metabolic measurements for marine animals spanning pelagic and benthic habitats across the full ocean depth gradient. Data were extracted from published laboratory, *in situ* and shipboard studies reporting oxygen consumption together with body mass, environmental temperature, sampling depth, and taxonomic identity [2, 6, 11, 12, 15, 21, 27, 28, 34, 36, 37, 39, 44–46, 48, 50, 51]. The final dataset comprised 689 metabolic measurements representing 11 marine phyla and 22 taxonomic classes, including fishes, crustaceans, molluscs, echinoderms, annelids, cnidarians, poriferans, and other major marine invertebrate groups. Metabolic rates reported were converted to a common unit of *µ*mol O_2_ h^−1^ using standard gas-volume conversions. All body masses are reported in grams wet weight. Environmental temperature was converted to Kelvin for all analyses. Hydrostatic pressure (atm) was calculated from sampling depth assuming a surface pressure of 1 atm and a hydrostatic increase of approximately 1 atm per 10.06 m depth. Hydrostatic pressure was not measured directly but calculated from sampling depth assuming standard seawater density. Because hydrostatic pressure is determined almost entirely by depth in the open ocean, this approximation introduces negligible error relative to other sources of variation in metabolic measurements.

To illustrate the predicted effects of hydrostatic pressure across the ocean water column, we used the annual mean global ocean temperature climatology from the World Ocean Atlas 2023 [33]. Temperature values were extracted at the standard depth levels and converted to Kelvin. A continuous temperature profile was obtained by linear interpolation between successive depth levels.

### B. Statistical Analysis

Linear models were fitted using ordinary least squares. Mixed-effects models were fitted with the lme4 package using maximum likelihood estimation for model comparison [4]. Competing models were compared using Akaike Information Criterion (AIC) and likelihood-ratio tests. Final parameter estimates were obtained using restricted maximum likelihood (REML).

